# ACE2 expression and localization are regulated by CFTR: implications beyond cystic fibrosis

**DOI:** 10.1101/2021.11.19.469220

**Authors:** Valentino Bezzerri, Valentina Gentili, Martina Api, Alessia Finotti, Chiara Papi, Anna Tamanini, Debora Olioso, Martina Duca, Erika Tedesco, Sara Leo, Monica Borgatti, Sonia Volpi, Paolo Pinton, Giulio Cabrini, Roberto Gambari, Francesco Blasi, Giuseppe Lippi, Alessandro Rimessi, Roberta Rizzo, Marco Cipolli

## Abstract

As an inherited disorder characterized by severe pulmonary disease, cystic fibrosis (CF) could be considered a comorbidity for coronavirus disease 2019 (COVID-19)^1^. Instead, CF seems to constitute an advantage in COVID-19 infection^2-5^.

To clarify whether host factors expressed by the CF epithelia may influence COVID-19 progression, we investigated the expression of SARS-CoV-2 receptor and coreceptors in primary airway epithelial cells. We found that angiotensin converting enzyme 2 (ACE2) expression and localization are regulated by cystic fibrosis transmembrane conductance regulator (CFTR) channels. Consistently, our results indicate that dysfunctional CFTR channels alter susceptibility to SARS-CoV-2 infection, resulting in reduced viral infection in CF cells. Depending on the pattern of ACE2 expression, the SARS-CoV-2 spike (S) protein induced high levels of Interleukin (IL)-6 in healthy donor-derived primary airway epithelial cells but a very weak response in primary CF cells. Collectively, these data support the hypothesis that CF condition is unfavorable for SARS-CoV-2 infection.

---

The genome of severe acute respiratory syndrome coronavirus 2 (SARS-CoV-2) encodes 28 proteins, including four structural proteins: spike (S), membrane, envelope and nucleocapsid. The S glycoprotein of SARS-CoV-2 is responsible for viral entry through binding of the ACE2 receptor^6-8^. Once the S protein has bound the ACE2 receptor, it is processed by several proteases, including transmembrane protease serine 2 (TMPRSS2) and furin, which promote priming of the S protein and the fusion of viral and cellular membranes^6^. SARS-CoV-2 infection primarily targets the airway epithelia. In some cases, infection leads to bilateral pneumonia with diffuse alveolar damage, which in turn may promote acute respiratory distress syndrome (ARDS), especially in subjects with comorbidities. Moreover, the onset of a cytokine storm, during which interleukin (IL)-6 plays a key role in morbidity, has been described in subcohorts of COVID-9 patients with poor prognosis^9,10^. CD13 is a transmembrane metalloprotease coregulated with ACE2 in various primate and human tissues^11^ that has also recently been proposed as a potential coreceptor of SARS-CoV-2 involved in viral entry and immune response^12^. Importantly, CD13 activation has been associated with the release of high levels of IL-6 in different cell models^13-15^.

CF is caused by mutations in the *CFTR* gene, which encodes a chloride and bicarbonate channel widely expressed in human epithelia. Loss of *CFTR* expression or function in airway epithelia is associated with reduced airway surface liquid (ASL) volume and dehydration. This process has been suggested as the initiating event of CF airway disease pathogenesis, which is characterized by severe impairment of lung function^16^. CF could therefore be considered an unfavorable comorbidity in patients with COVID-19, particularly considering that other respiratory viral infections, including respiratory syncytial virus (RSV) and influenza A (H1N1), lead to a rapid deterioration of lung function and increased mortality in CF patients^17,18^.

However, several studies conducted on Belgian^2^, French^5^, Spanish^4^ and Italian^19^ cohorts of CF patients have reported that CF seems to protect against COVID-19. Even though age distribution has been proposed as a major confounding factor for incidence calculation in these studies^20^, the European Society of Cystic Fibrosis (ECFS) recently concluded that the case fatality rate associated with COVID-19 in CF patients was lower than in the general population. On the other hand, considering data that emerged from the French and Italian cohorts, it should not be dismissed that post lung transplant patients exhibited more severe manifestations in response to COVID-19^5,19^.

Since 51.2% of patients registered in the ECFS patient registry are adults^20^, the possibility that the favorable COVID-19 disease outcome could be dependent only on pediatric age is questionable.

Therefore, our hypothesis was that some specific host factors associated with CF may influence susceptibility to SARS-CoV-2 infection. Interestingly, it has been recently reported that TMEM16F, a Ca^2+^-activated chloride channel, plays a key role in SARS-CoV-2 viral entry and syncytia formation in lung epithelial cells^21^, may be actively involved in the pathogenesis of COVID-19. Furthermore, it has already been established that CFTR can regulate other apical proteins, including the ion channel solute carrier family 26 member 9 (SLC26A9)^22^, the epithelial sodium channel (ENaC)^23^, the potassium channel KIR 1.1^24^, Phosphatase And Tensin Homolog protein (PTEN)^25^, and receptors such as the A2B adenosine receptor^26^. These regulatory functions of CFTR might be due to direct binding of CFTR to other proteins or indirect binding through PDZ-interacting domains through the adapter proteins ezrin, Na^+^/H^+^ exchanger regulatory factor (NHERF)1 and NHERF2. Given all of these possibilities, we investigated the role of CFTR in regulating SARS-CoV-2 receptors and coreceptors.

We first explored the expression of *ACE2, TMRPSS2*, and *CD13* in CF primary airway epithelial cells. *ACE2* mRNA expression was reduced in both primary bronchial epithelial cells (HBECs) and nasal epithelial cells (HNECs) isolated from CF patients homozygous for the F508del *CFTR* mutation (Supplementary Table 1) compared to healthy donor-derived tissues (Fig. 1a). Consistent with the mRNA data, ACE2 and CD13 protein levels were remarkably lower in primary well-differentiated CF-HNECs (Fig. 1b). In contrast, no change in *TMPRSS2* expression was observed between CF and nonCF primary HNECs, either in terms of mRNA or protein (Fig. 1a and b). To determine whether *CFTR* expression influences *ACE2* expression, we utilized different cell models in which gene editing approaches were applied to modulate *CFTR* expression. In particular, we employed the human lung adenocarcinoma Calu-3 cell line in which *CFTR* was stably silenced by a short hairpin (sh)RNA (SH3 cells), as previously described^27^, the well-established CF bronchial epithelial cell line CFBE41o-, in which *CFTR* expression is almost undetectable, and cell lines derived from CFBE41o- cells expressing wild-type *CFTR* (CFBE41o-WT) or F508del-*CFTR* (CFBE41o-F508del). Finally, we used a human bronchial epithelial 16HBE14o-cell line expressing wild-type *CFTR* and edited using the CRISPR/Cas9 approach to obtain two supplemental cell lines carrying biallelic W1282X-*CFTR* or G542X-*CFTR* nonsense mutations.

**Fig. 1.**
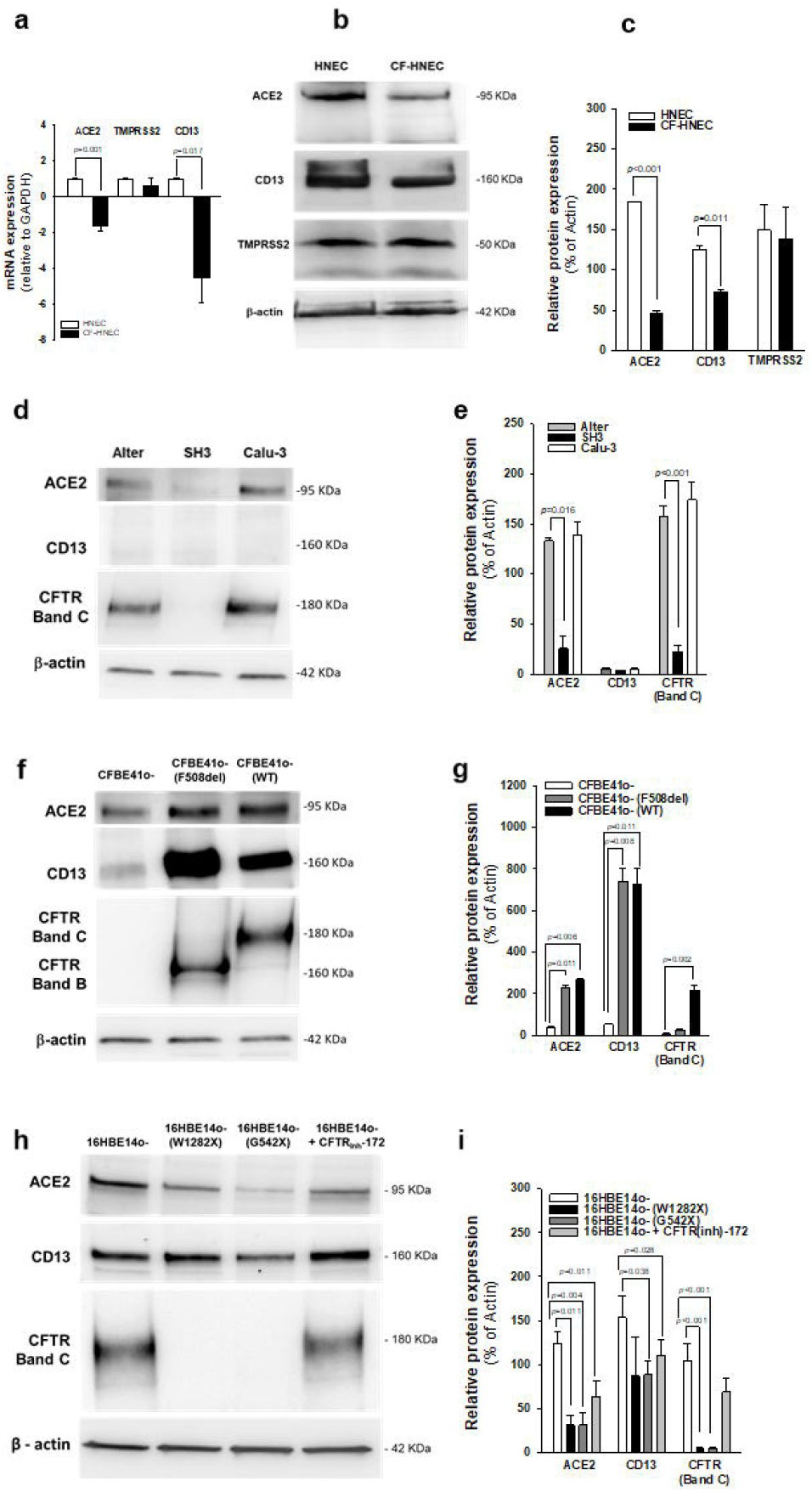
ACE2 and CD13 expression is reduced in CF. **a-b**, Quantification of ACE2, TMPRSS2 and CD13 mRNA expression by qPCR in lysates collected from well-differentiated HNECs grown at the air-liquid interface derived from a pool of 14 healthy donors (HNECs) and 2 CF patients (CF-HNECs) (**a**). The genotypes of the patients enrolled in this study are reported in Supplementary Table 1. Data are shown as the mean ± SEM of four independent experiments performed in duplicate. Normal distribution was confirmed using the Shapiro–Wilk test before running Student’s t-test for paired data, which has been reported in the histograms. **b**, Representative western blot analysis of protein extracts from differentiated HNECs grown at the air-liquid interface obtained from a pool of 14 healthy control subjects versus CF-HNECs obtained from an F508del-CFTR homozygous CF patient. Densitometry analysis (% of β-actin expression) of three independent experiments, conducted as reported in b, is depicted. **c. d**, western blot analysis of ACE2, CD13 and CFTR and densitometry analysis (% of β-actin expression) of three independent experiments (**e**) performed in protein extracts from Calu-3, SH3 and Alter, 16HBE14o- and CFBE41o-expressing F508del-CFTR (F508del) or wild-type CFTR (WT) (**f-g**), and 16HBE14o- cells carrying biallelic W1282X-CFTR or G542X-CFTR mutations, or incubated with 5 μM CFTR inhibitor CFTR(inh)-172 for 24 h (**h-i**).

The results unambiguously indicated that *CFTR* expression is associated with *ACE2* expression (Fig. 1). In particular, *CFTR*-deficient SH3 cells displayed a remarkable reduction in ACE2 levels compared to both parental Calu-3 cells and mock-transfected cells, i.e. Alter^27^ cells (Fig. 1d and e). CD13 is not expressed in these cells. In contrast, the introduction of both mutated F508del-*CFTR* and wild-type *CFTR* in CFBE41o- cells resulted in a substantial increase in ACE2 levels, accompanied by remarkable induction of CD13 protein expression (Fig. 1f and g). Then we assessed *ACE2* expression in 16HBE14o- cells in which the *CFTR* sequence was edited using CRISPR/Cas9 technology. Consistent with the results from SH3 and Alter cells, we found that ACE2 protein synthesis was remarkably reduced in cells harboring nonsense-mutated *CFTR*, with undetectable levels of CFTR protein (Fig. 1h and i). Thus, it seems that variations in ACE2 expression are not associated with specific *CFTR* genetic classes of defect, such as Class II folding defect of F508del-CFTR, but rather, are more widely related to defective CFTR protein. Moreover, a well-established inhibitor of CFTR ion flux, thiazolidinone CFTR(inh)-172^28^, partially reduced the expression of ACE2 (Fig. 1h and i). Once again, CD13 protein levels were partially corepressed in these cellular models, particularly in the G542X-*CFTR* cell line (Fig. 1h and i), confirming that CD13 expression is coregulated with ACE2 expression in airway epithelial cells, as previously demonstrated in other cell types^11^.

We therefore decided to investigate whether the subcellular distribution of the ACE2 receptor is dependent on CFTR expression, localization and function considering that the CFTR channel exists in a multiprotein complex that regulates CFTR channel activity but may also regulate the localization and function of other proteins on the plasma membrane^23-26^. For this purpose, we investigated the subcellular distribution of ACE2 in different CFBE41o-cell models using immunofluorescence. In CFBE41o- cells, both CFTR and the ACE2 receptor were not localized to the plasma membrane (Fig. 2a). The typical plasma membrane localization of the ACE2 receptor was instead observed in CFBE41o-(WT) cells expressing wild-type *CFTR* (Fig. 2a). The expression of wild-type CFTR protein on the plasma membrane was strictly associated with the subcellular distribution of ACE2, as confirmed by Manders M1 and Pearson’s coefficients, parameters sustaining the correlation between the intracellular localization of CFTR and ACE2 (Fig. 2b). Interestingly, in CFBE41o- cells expressing F508del-*CFTR*, which encodes an unfolded CFTR protein that is primarily retained in the ER, ACE2 was almost entirely localized to the ER (Fig. 2a). Once again, the colocalization of ACE2 with CFTR was validated by Manders M1 and Pearson’s coefficients (Fig. 2b). The redistribution of ACE2 to the ER was confirmed by subcellular quantification, since the ratio between plasma membrane fluorescence and ER fluorescence was significantly reduced in CFBE41o-(F508del) cells compared to CFBE41o-(WT) and 16HBE14o- cells, where both CFTR and ACE2 proteins were primarily localized on the plasma membrane (Fig. 2b, III). Next, we examined whether *ACE2* expression and localization could be only influenced by *CFTR* expression, or whether CFTR channel function could play a role. To this aim, we incubated 16HBE14o- and CFBE41o-(WT) cells re-expressing wild-type *CFTR* with CFTR(inh)-172 for 24 hours. Our data indicated that the prolonged functional block of ion efflux from CFTR does not lead to modification of the subcellular localization of ACE2 (Fig. 2a and b).

**Fig. 2.**
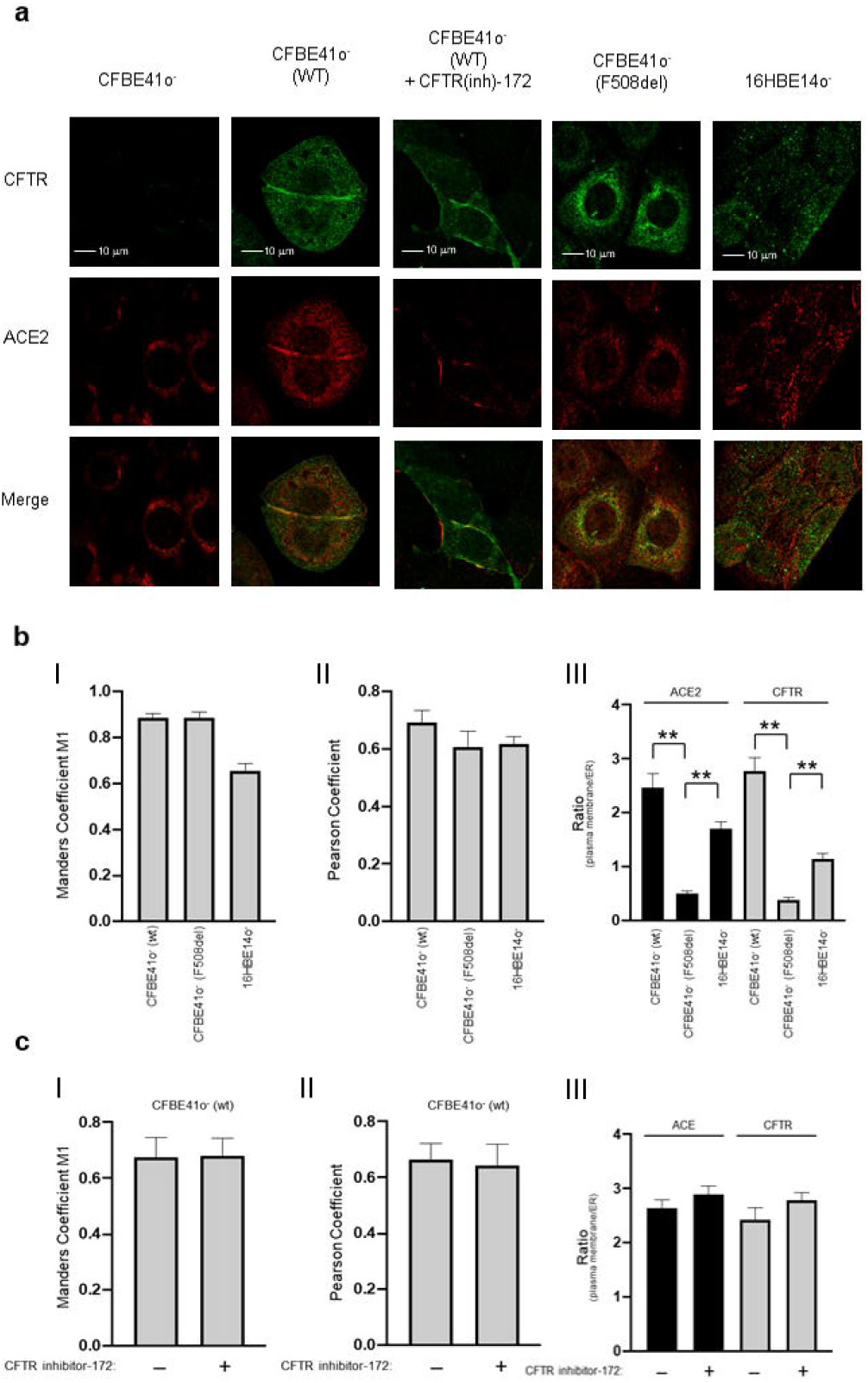
Localization of CFTR on the plasma membrane, but not its function, is essential for the subcellular localization of ACE2 on the cell surface. **a)** Representative images of immunofluorescence detection of CFTR (green) and ACE2 receptor (red) under basal conditions in CFBE41o- and CFBE41o-expressing wild-type CFTR (before and after 24 hours of 5 μM CFTR inhibitor-172 treatment), CFBE41o-expressing F508del-CFTR and 16HBE14o- cells. Images were acquired with a Zeiss LSM510 confocal microscope (scale bar: 10 µm). **b)** Quantification of colocalization between CFTR and the ACE2 receptor and quantification of the subcellular distribution of ACE2 and CFTR. Manders M1 **(I)** and Pearson’s **(II)** coefficients represent the correlation between the intracellular localization of CFTR and ACE2 in the different cell lines. **(III)** Quantification of the subcellular distribution of the ACE2 receptor (black) and CFTR (gray) in the different cell models. The histograms represent the ratio between plasma membrane fluorescence and endoplasmic reticulum (ER) fluorescence. **c)** Quantification of colocalization between CFTR and ACE2 in CFBE41o-expressing wild-type CFTR (WT) in the presence (+) or absence (-) of the CFTR inhibitor-172. The histograms show the Manders M1 **(I)** and Pearson’s **(II)** coefficients under different experimental conditions. **(III)** The histograms represent the ratio between plasma membrane fluorescence and endoplasmic reticulum (ER) fluorescence of ACE2 (black) and CFTR (gray). Data are mean ± SEM of 6 independent experiments. Normal distribution was tested by the Shapiro–Wilk test before running the Student’s t test for paired data, which has been reported within the histograms (**p<0.01).

To exclude the possibility that mislocalization of the ACE2 receptor into the ER is due to alterations in the membrane trafficking pathway associated with the unfolded CFTR protein, we investigated the subcellular localization of CFTR-interactor NHERF1 ^29^. In the absence of *CFTR* expression, NHERF1 localizes closely to the plasma membrane in CFBE41o- cells (Fig. 3a). NHERF1 normally binds to the CFTR channel through its PDZ domains. This interaction was indeed confirmed by the coefficients of colocalization in CFBE41o-(WT) cells re-expressing wild-type *CFTR* (Fig. 3b). However, the typical localization of NHERF1 was perturbed in CFBE41o- cells expressing the mutated F508del *CFTR*, encoding a CFTR protein that is primarily retained within the ER (Fig. 3b). These results therefore support the hypothesis that the presence of a mutated CFTR channel, as well as the loss of *CFTR* expression, may induce mislocalization of the ACE2 receptor regardless of the membrane trafficking pathways.

**Fig. 3.**
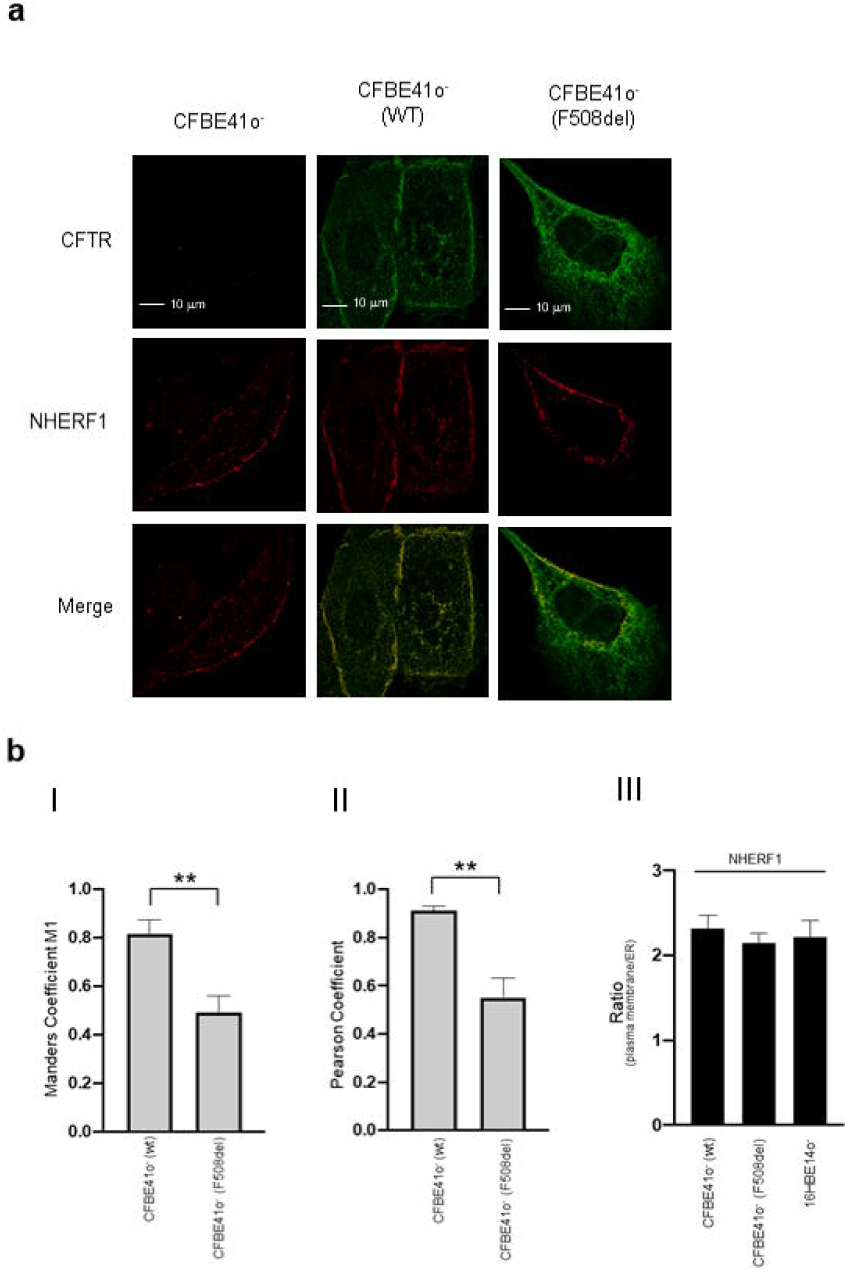
CFTR subcellular localization does not affect localization of the NHERF1 scaffolding protein on the plasma membrane. **a)** Representative images of immunofluorescence detection of CFTR (green) and NHERF1 (red) under basal conditions in CFBE41o-, CFBE41o-expressing CFTR, CFBE41o-expressing and F508del-CFTR (scale bar: 10 µm). **b)** Quantification of colocalization between CFTR and NHERF1 and subcellular distribution of EPB50 in different cell lines. Manders M1 **(I)** and Pearson’s **(II)** coefficients represent the correlation between the intracellular localization of CFTR and NHERF1, whereas **(III)** represents the ratio between plasma membrane fluorescence and endoplasmic reticulum (ER) fluorescence of NHERF1. Data are mean ± SEM of 7 independent experiments. Normal distribution was tested by the Shapiro–Wilk test before running the Student’s t test for paired data, which has been reported within the histograms (**p<0.01).

In contrast to what has been observed for ACE2, the plasma membrane localization of CD13 was not perturbed by *CFTR* expression (Fig. 4).

**Fig. 4.**
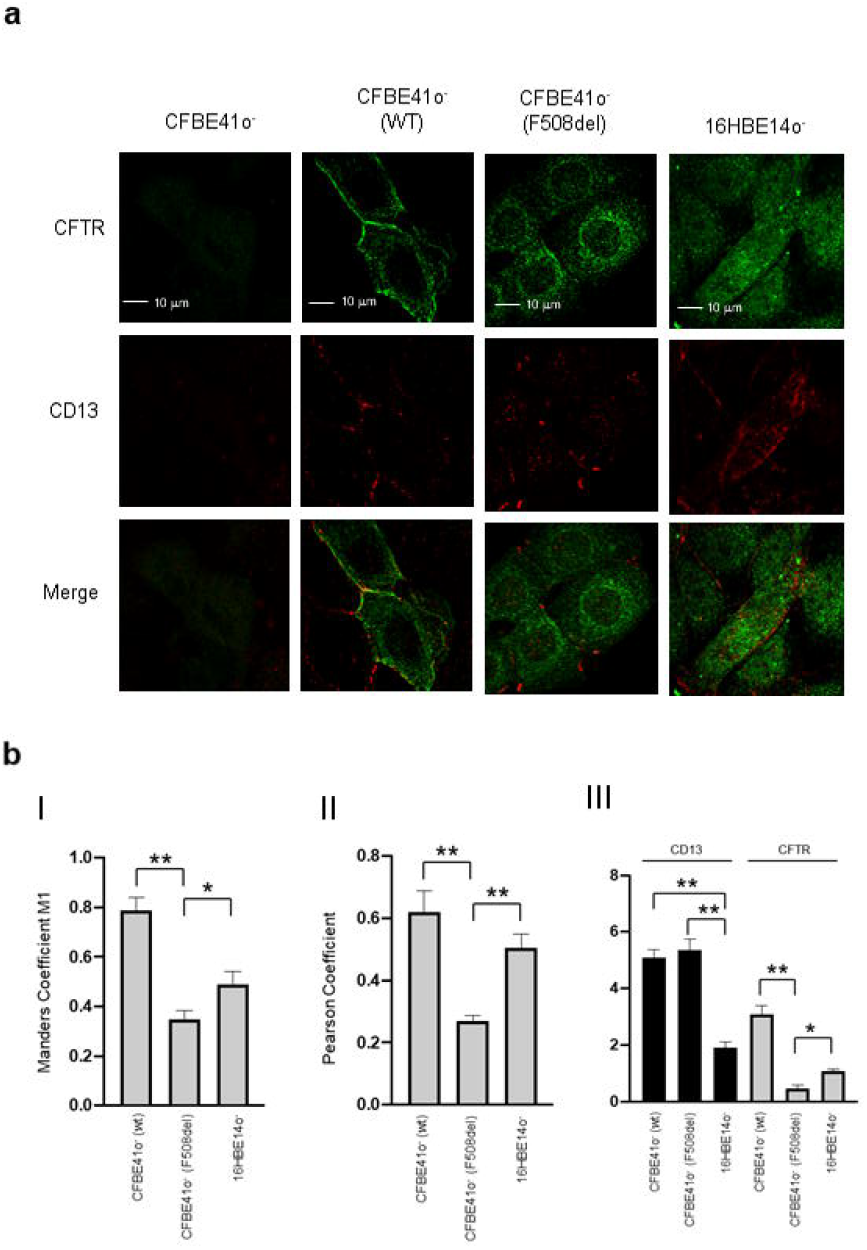
CFTR expression controls expression of CD13 without affecting its localization. **a)** Representative images of immunofluorescence detection of CFTR (green) and CD13 receptor (red) under basal conditions, acquired by Zeiss LSM510 confocal microscopy, in CFBE41o-, CFBE41o-expressing wild-type CFTR (WT), CFBE41o-expressing F508del-CFTR and 16HBE14o-are shown (scale bar: 10 µm). **b)** Colocalization analysis between CFTR and CD13 is represented by the Manders M1 **(I)** and Pearson’s **(II)** coefficients shown in the figure for the different cell models, whereas **(III)** represents the ratio between plasma membrane fluorescence and endoplasmic reticulum (ER) fluorescence of CD13 (black) and CFTR (gray). Data are mean ± SEM of 6 independent experiments. Normal distribution was tested by the Shapiro–Wilk test before running the Student’s t test for paired data, which has been reported within the histograms (*p<0.05, **p<0.01)

To confirm the role of CFTR in the SARS-CoV-2 life cycle, we employed two complementary strategies, use of a pharmacological inhibitor of CFTR function and a miRNA-based approach to specifically downregulate CFTR production, using the CFBE41o- and Calu-3 cell lines. The chronic effect of CFTR(inh)-172 on the extracellular release of SARS-CoV-2 from infected CFBE41o-(WT) cells expressing wild-type *CFTR* is shown in Fig. 5a. We utilized a previously validated methodology for in vitro viral infections^30^. The data obtained support the concept that inhibition of CFTR function is associated with a remarkable reduction in SARS-CoV-2 virulence (Fig. 5a). For comparison, the replication of SARS-CoV-2 has also been investigated in the mutant CFBE41o-(F508del) and parental CFBE41o-cell lines. Using these two cell lines, which harbor nonfunctional CFTR channels, the replication of SARS-CoV-2 was strongly suppressed (Fig. 5a).

**Fig. 5.**
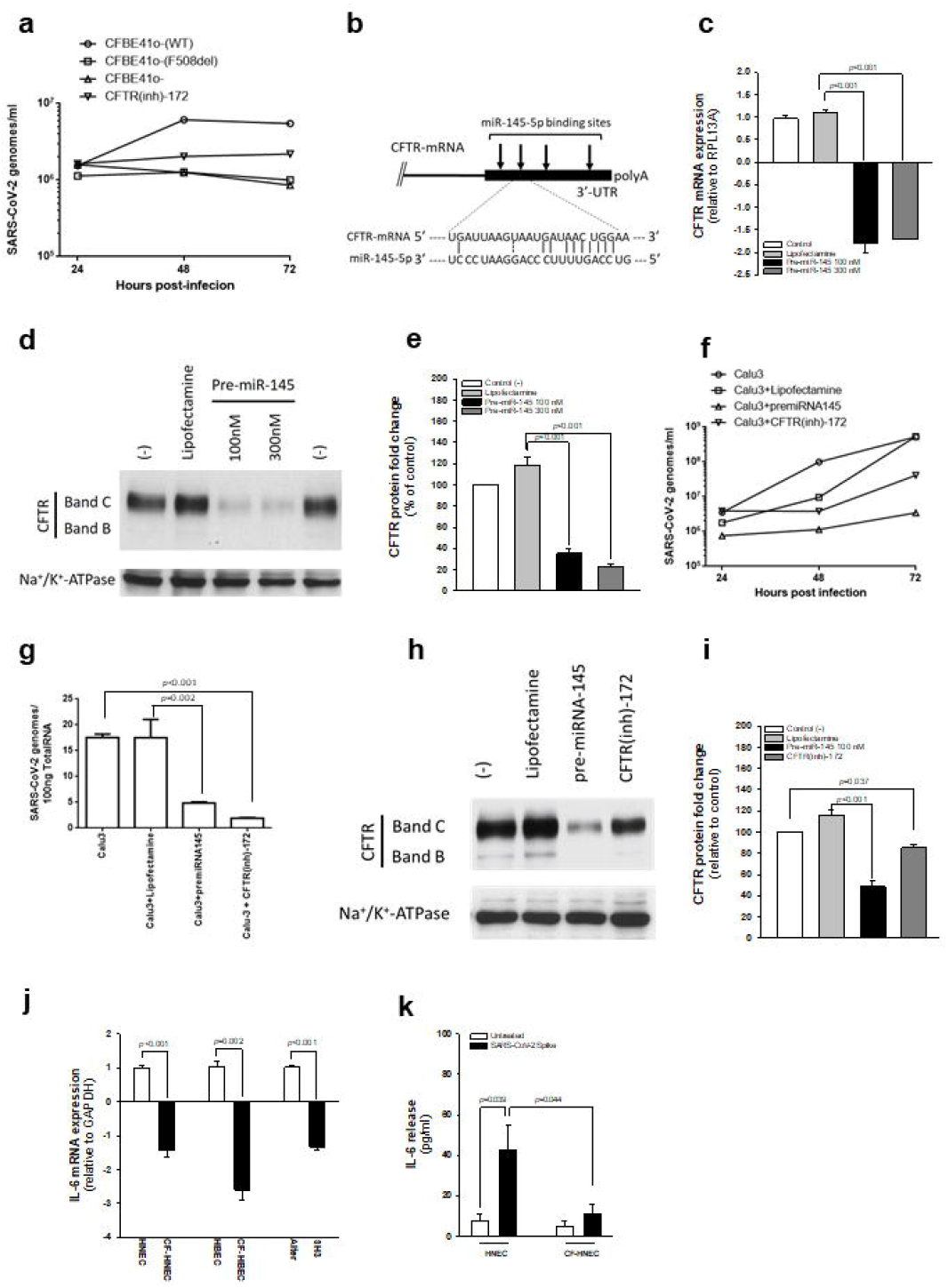
Loss of CFTR expression and function inhibits SARS-CoV-2 entry and replication and reduces spike-dependent IL-6 release in airway epithelial cells. **a**, CFBE41o-(F508del) or CFBE41o-(WT) (before and after 5 μM CFTR(inh)-172 treatment for 24h) were infected with 0.1 MOI SARS-CoV-2 for 1 hour, and then the viral titer in cell supernatants was evaluated 24, 48 and 72 hours post infection by RT– qPCR. The results are shown as the mean ± SEM from three independent experiments. **b**, Location of the miR-145-5p binding sites within the CFTR 3’-UTR (binding site displaying the highest affinity is shown). **c-e**, Effect of premiR-145-5p treatment in Calu-3 cells. The effect on CFTR mRNA was verified by RT–qPCR **(c)**. The effects on CFTR content were evaluated by western blot analysis **(d). e**, Densitometry analysis of four independent experiments, conducted as reported in d. **f-g**, Effect of premiR-145-5p compared to CFTR(inh)-172 on the extracellular release of SARS-CoV-2 in infected Calu-3 cells. Extracellular release of SARS-CoV-2 genomes **(f)**. Results are reported as the mean ± SD from three independent experiments. Intracellular production of SARS-CoV-2 genomes **(g)**. Results are reported as the mean ± SD from three independent experiments. **h-i**, Western blot analysis of CFTR in response to SARS-CoV-2 infection under the same conditions depicted in (f-g). **j-k**, Constitutive IL-6 expression is reduced in CF airway epithelial cells, and IL-6 release upon SARS-CoV-2 spike protein stimulation is reduced in CF. **j**, IL-6 mRNA was quantified by RT–qPCR in lysates obtained from differentiated HNEC cells obtained from a pool of 14 healthy donors (HNEC) versus 2 CF patients (CF-HNEC) homozygous for F508del *CFTR* or in primary HBEC obtained from four healthy control subjects versus four CF patients homozygous for F508del *CFTR* or in Alter cells versus SH3 cells. **k**, Quantification of IL-6 release by ELISA in cell culture supernatants obtained from well-differentiated HNECs and CF-HNECs. Cells were stimulated on both apical and basolateral sides with 1 μg/ml recombinant SARS-CoV-2 S protein for 12 h. Data are reported as the mean ± SEM from three independent experiments performed in duplicate. Student’s t-test is reported in the histograms.

Regarding the miRNA-based strategy, our group and others have previously demonstrated that *CFTR* expression is under posttranscriptional control of different microRNAs^31-34^. Since *CFTR* expression increased in response to antagomiR molecules against miR-145-5p^31,32,35^, the exposure of bronchial epithelial cells to this miRNA might lead to *CFTR* downregulation (see Fig. 5b for localization of miR-145-5p binding sites within the CFTR 3’UTR). In fact, four miR-145-5p binding sites are present in the 3’-UTR of *CFTR* (Fig. 5b). Treatment of Calu-3 cells with premiR-145-5p significantly reduced the accumulation of *CFTR* mRNA (Fig. 5c). Most importantly, treatment of Calu-3 cells with premiR145-5p remarkably reduced CFTR protein levels (Fig. 5d and e). The conclusion of these experiments fully supports the concept that downregulation of *CFTR* can be achieved by a miRNA-mimicking strategy using premiR-145-5p.

We next assessed whether this treatment, performed on SARS-CoV-2-infected Calu-3 cells, is associated with alteration of the SARS-CoV-2 life cycle. The results revealed that pretreatment of Calu-3 cells with 100 nM premiR-145-5p is sufficient to dramatically inhibit SARS-CoV-2 viral production in a similar fashion to that observed by incubating Calu-3 cells with the CFTR(inh)-172 compound (Fig. 5f and g). We observed partial inhibition of mature CFTR protein expression upon prolonged incubation (72h) of CFTR(inh)-172 (Fig. 5h and i). This phenomenon has already been reported in other human cell lines, including K562 and SUP-B15^36^, and might explain the partial reduction of ACE2 observed upon CFTR(inh)-172 incubation shown in Fig. 1. As expected from the data shown in Fig. 5c-e, western blot analysis of CFTR protein content in response to the different treatment conditions confirmed that CFTR synthesis was greatly inhibited by treatment with premiR145-5p (Fig. 5h and i).

In addition, we evaluated the effect of the ACE2 ligand SARS-CoV-2 S protein on IL-6 expression. Although CF lung pathology has been established as a proinflammatory condition featuring elevated levels of cytokines and chemokines, in particular IL-8^37^, some clinical evidence suggests that IL-6 levels are extremely low in sputum samples and bronchoalveolar lavage fluids obtained from CF patients^38-40^. Our data consistently showed that IL-6 mRNA expression was constitutively reduced in primary nasal and bronchial epithelial cells obtained from CF patients compared to healthy donor-derived tissues (Fig. 5j). Interestingly, knockdown of *CFTR* expression in SH3 cells reduced expression of *IL6* mRNA compared to mock-transfected Alter cells (Fig. 5j). Most importantly, stimulation with the SARS-CoV-2 S protein led to a remarkable induction of IL-6 release in well-differentiated primary HNECs grown at the air-liquid interface derived from healthy donors but induced a very weak response in CF HNECs (Fig. 5k). These data are in line with a recent report showing that the SARS-CoV-2 S protein increases IL-6 expression in macrophages isolated from nonCF subjects but not in CF cells^41^, strengthening the observations of the reduced expression and mislocalization of the ACE2 receptor in CF cells.

In conclusion, this work provides new insights into regulation of the expression and localization of ACE2 in relation to CFTR expression and function. This issue is of substantial interest for unraveling additional aspects in the pathogenesis of SARS-CoV-2 infection. Here, we show that CFTR expression and function are directly correlated with *ACE2* expression and their interaction is necessary to place ACE2 receptor on the plasma membrane. Of note, our results show that CF airway epithelial cells constitutively express mislocalized and lower levels of ACE2 receptor than nonCF cells, which in turn leads to a reduction in SARS-CoV-2 infectivity. These results are in line with a previous report on TMEM16F^21^, strengthening the hypothesis that chloride channels are considerably involved in the SARS-CoV-2 biological cycle.

Given these findings, the use of chloride channel inhibitors or miRNAs against *CFTR*, such as miR-145-5p, is conceivable as a possible therapeutic strategy for preventing or limiting the adverse consequences of SARS-CoV-2 infection in humans.

## Methods

### Human samples

All human samples were obtained and analyzed in accordance with the Declaration of Helsinki after written consent was obtained. All protocols were approved by the Ethics Committee of the Azienda Ospedaliera Universitaria Integrata (Verona, Italy), approval No. 2917CESC. Well-differentiated Mucil-air® primary human nasal epithelial cells from F508del/F508del CF patients (CF-HNEC) or from healthy donors (HNEC) were supplied by Epithelix (Plan-les-Ouates, Switzerland) after strict quality control was performed by the supplier. HNECs were cultured on Snapwell supports with Mucil-air® differentiating medium (Epithelix) as previously described^42^. Cell lysates from primary human bronchial epithelial cells from F508del/F508del CF patients (CF-HBECs) or from healthy donors (HBECs) were supplied by Epithelix. The human samples analyzed in this study are summarized in supplementary Table 1.

### Cell culture

CF bronchial epithelial CFBE41o- cells with or without stable expression of F508del-CFTR or wild-type CFTR obtained by Dr. J.P. Clancy^43^ cells were cultured in MEM supplemented with 10% fetal bovine serum (FBS) and 2 mM L-glutamine in the absence of antibiotics. The submucosal gland cell line Calu-3, generated from bronchial adenocarcinoma, together with SH3 and Alter cells were kindly provided by Dr M. Chanson^27^. Briefly, stable expression of short hairpin RNAs (shRNAs) against CFTR was induced in Calu-3 cells by transfecting the Sleeping Beauty transposon vector pT2/si-PuroV2, generating a CFTR knockout cell line (SH3). A scrambled shRNA sequence was used to generate a mock transfected cell line (Alter), as previously described^27^. Calu-3, SH3, and Alter cells were maintained in DMEM/F12 (3:1 vol/vol) supplemented with 10% FBS without streptomycin or penicillin but were continuously selected in the presence of 4 μg/ml puromycin, as previously described^27^.

#### Inhibition of CFTR expression by pre-miR-145

Transfection procedure of pre-miR-145 in Calu-3 cells was performed in 12-well plates using Lipofectamine RNAiMAX Transfection Reagent (Invitrogen, Thermo Fischer Scientific) accordingly to manufacturer’s instruction. Calu-3 cells were seeded at 2.5×10^5^/ml with 100 or 300 nM of hsa-miR-145-5p miRNA precursor (PM11480, Ambion, Thermo Fisher Scientific). After 72 h, cells were collected and total RNA was isolated using TRI Reagent™ (Sigma Aldrich) and immediately converted to cDNA. For the experiments with SARS-CoV-2 infection, cells were pre-treated with pre-miR-145-5p at 100nM for 48 hours before the infection.

### Quantitative PCR

Total RNA from HNECs, HBECs, Calu-3, SH3, Alter, CFBE41o- and 16HBE14o- cells was isolated using a High Pure RNA Isolation Kit (Roche, Mannheim, Germany) following the manufacturer’s protocol. RNA concentration was determined using a NanoDrop 2000 spectrophotometer (Thermo Fisher Scientific, Waltham, MA) and then stored at -80°C until use. A total of 500 ng of RNA was reverse transcribed to cDNA using a High-Capacity cDNA Reverse Transcription Kit with random primers (Thermo Fisher Scientific) following the manufacturer’s recommendations. A total of 25 ng of cDNA was used for each reaction to quantify the relative gene expression. cDNA was then amplified using qPCRBio SyGreen Mix (PCR Biosystems, Wayne, PA, USA) and QuantiTect Primer Assays (Qiagen, Hilden, Germany) for ACE2 (Hs_ACE2_1_SG, NM_021804), TMPRSS2 (Hs_TMPRSS2_1_SG, NM_005656), CD13/ANPEP (Hs_ANPEP_1_SG, NM_001150), and GAPDH (HS_GAPDH_1_SG, NM_001256799). The analysis was performed using an AriaMx thermocycler (Agilent, Santa Clara, CA). Changes in mRNA expression levels were calculated following normalization to the GAPDH reference gene, and relative quantification was performed using the comparative cycle threshold method.

In experiments employing the pre-miR-145, CFTR expression was analyzed by RT-qPCR using 300 ng of total RNA, which were reverse transcribed using the Taq-Man Reverse Transcription PCR Kit and random hexamers (Applied Biosystems, Thermo Fischer Scientific) in a final reaction volume of 50 µl. CFTR gene-specific double fluorescently labeled probe and primers were used (Assay ID: Hs00357011_m1). The relative expression was calculated using the comparative cycle threshold method and, as reference gene, the human RPL13A (Assay ID: Hs03043885_g1). Assays were purchased from Applied Biosystems.

### Western blot

A total of 40□μg of cell extracts obtained from 16HBE14o-, CFBE41o-, Calu-3, SH3, Alter and primary HNEC and HBEC were denatured for 5□min at 95°C in 4x Laemmli Sample Buffer, (277.8 mM Tris-HCl, pH 6.8, 44.4% glycerol, 4.4% LDS, 0.02% bromophenol blue) (Bio–Rad Laboratoires, Philadelphia, PA), containing 355 mM 2-mercaptoethanol. For ACE2, CD13, TMPRSS2 and β-actin analysis, protein extracts were loaded on Miniprotean TGX (4-14%) SDS–PAGE gel (Bio–Rad Laboratories) in Tris-glycine buffer (25 mM Tris, 192 mM glycine, and 0.1% SDS) using Precision Plus dual color tag protein ladder (Bio–Rad Laboratories) to determine molecular weight. For CFTR analysis, protein extracts were loaded on Criterion XT (Bio–Rad laboratories) precast gels (3-8%) in XT Tricine buffer (Bio–Rad Laboratories) using Precision Plus Protein Kaleidoscope standards (Bio–Rad Laboratories). Gels were then transferred onto Transblot turbo PVDF membranes (Bio–Rad laboratories) at 20 V using Transblot Turbo equipment (Bio–Rad laboratories) for 5-12 minutes. Membranes were blocked in 5% BSA (ACE2, TMPRSS2, CD13, β-actin) or 5% nonfat dried milk (PanReac AppliChem, Darmstadt, Germany) for 90 minutes at room temperature and washed with TBS (5 mM Tris-HCl pH 7.6, 150□mM NaCl) supplemented with 0.05% Tween-20 (TBS/T) for 15 minutes. Membranes were probed with primary anti-human ACE2 rabbit polyclonal antibody (ab15348, Abcam, Cambridge, UK), anti-TMPRSS2 rabbit monoclonal antibody (ab109131, Abcam), or anti-CD13 rabbit monoclonal antibody (ab108382, Abcam) in TBS/T with 5% BSA or mouse IgG_2b_ anti-CFTR NBD2 (antibody 596, Cystic Fibrosis North American Foundation, Chapel Hill, NC, 1:2500 dilution) in 5% nonfat dried milk and incubated overnight at 4°C. After washing, membranes were incubated with mouse anti-rabbit IgG-horseradish peroxidase-coupled secondary antibody (sc2357, Santa Cruz Biotechnology, Dallas, TX, USA, dilution 1:2000) or donkey anti-mouse IgG-horseradish peroxidase-coupled secondary antibody (RnD Systems) diluted in TBS/T for 90 minutes at room temperature. Chemiluminescence was measured using Clarity Western ECL Substrate (Bio–Rad Laboratories), incubating membranes in ECL for 5□min at room temperature. Band intensity was evaluated by scanning video densitometry using a ChemiDoc imaging system (UVP, LCC, Upland, CA). Blots were then reprobed with monoclonal anti-β-actin-peroxidase clone AC-15 antibody (a3854, Merck, Saint Louis, MO, dilution 1:5000) in TBS/T for 90 minutes. For western blotting, cell pellets were lysed in 1% Nonidet P40 (IGEPAL), 0.5% sodium deoxycholate, 200 mM NaCl, 10 mM Trizma base, pH 7.8, 1 mM EDTA plus protease inhibitor mixture and 1 mM PMSF for 30 min on ice. Lysates were cleared by centrifugation at 10,000× g for 10 min at 4°C. Protein concentration was determined using the Lowry method after precipitation with 5% trichloroacetic acid (TCA), utilizing bovine serum albumin (BSA) as a standard. For CFTR analysis, 20 μg of the total proteic extracts were heated in Laemmli buffer (Bio–Rad, Hercules, CA, USA) at 37°C for 10 min and loaded onto a 3 to 8% Tris-acetate gel (Bio–Rad, Hercules, CA, USA). Proteins were transferred to polyvinylidene difluoride (PVDF) membranes (Bio–Rad, Hercules, CA, USA) using Trans Blot Turbo (Bio–Rad Laboratories) and processed for western blotting using a mouse monoclonal antibody, clone 596, against the NBD2 domain of CFTR (University of North Carolina, Cystic Fibrosis Center, Chapel Hill, NC, USA) at a dilution of 1:2500 with overnight incubation at 4°C. After washing, the membranes were incubated with horseradish peroxidase-coupled anti-mouse immunoglobulin (R&D System, Minneapolis, MN, USA) at room temperature for 1 h, and after washing, the signal was developed by enhanced chemiluminescence (LumiGlo Reagent and Peroxide, Cell Signaling, Danvers, MA, USA). After membrane stripping, a β-actin monoclonal antibody (Sigma Aldrich, St. Louis, MO, USA) was used to ensure equal loading of samples. For experiments employing the pre-miR-145, cellular extracts were obtained using Pierce RIPA Buffer (Thermo Fisher Scientific, Waltham, MA, USA) and then sonicated for 3×30 sec on ice at 50% amplitude using the Vibra-Cell VC130 Ultrasonic Processor (Sonics). Protein extracts were quantified using BCA Protein Assay kit (Thermo Fisher Scientific). Sample was heated 37 °C for 10 min in 1× sodium dodecyl sulfate (SDS) sample buffer (62.5 mM Tris–HCl pH 6.8, 2% SDS, 50 mM dithiothreithol (DTT) 0.01% bromphenol blue, 10% glycerol), and loaded on 7 % SDS– polyacrylamide gel in Tris–glycine buffer (25 mM Tris, 192 mM glycine, 0.1% SDS). Spectra Multicolore Broad Range Protein Ladder (Thermo Fisher Scientific) was used as standard to determine the molecular weight. Gels were transferred to nitrocellulose membrane (Thermo Fisher Scientific) using Trans Blot Turbo (Bio-Rad Laboratories). For CFTR analysis, the filter was processed using the mouse monoclonal antibody, clone 596, against the NBD2 domain of CFTR (University of North Carolina, Cystic Fibrosis Center, Chapel Hill, NC, US) at a dilution of 1:2500 by an overnight incubation at 4 °C. After washes, the membranes were incubated with horseradish peroxidase-coupled anti-mouse immunoglobulin (R&D System, Minneapolis, MN, USA) at room temperature for 1 h and after subsequent washes, the signal was developed by enhanced chemiluminescence (LumiGlo Reagent and Peroxide, Cell Signaling). The Na^+^/K^+^ ATPase protein (SC-514614, Santa Cruz Biotechnology) was used to confirm the equal loading of samples.

### Immunofluorescence

Cells were seeded onto 24 mm coverslips. The next day, the cells were rinsed with ice-cold PBS and fixed in 4% paraformaldehyde for 20 min at room temperature. To eliminate paraformaldehyde autofluorescence, cells were incubated for 10 min at room temperature with a 0.1 M glycine solution, washed and permeabilized with 0.1% Triton X-100. Nonspecific binding of antibodies was prevented by incubating cells with a 2% BSA solution for an hour at room temperature.

Subsequently, a double immunofluorescence procedure was performed, and the cells were incubated overnight at 4°C with a mouse anti-CFTR (antibody 570, cod. A2 from CF American Foundation, 1:200) and a rabbit anti-ACE2 (ab15348 from AbCam, 1:200) or anti-CD13 (ab108382 from AbCam, 1:200), or in a sequential procedure, with mouse anti-CFTR and mouse anti-NHERF1 (611161 from BD Transduction Laboratories, 1:200). Cells were then washed three times with cold PBS and incubated for 1 hour at room temperature with goat anti-mouse Alexa-488 (A32723 from Thermo Fisher Scientific, 1:1000) and goat anti-rabbit Alexa-594 (A11012 from Thermo Fisher Scientific, 1:1000) or, in the second case, with goat anti-mouse Alexa-488 (1:1000) and rabbit anti-mouse Alexa-594 (A27027 from Thermo Fisher Scientific, 1:1000) antibodies. Cells were then mounted on a coverslip with Pro-Long Gold antifade reagent (Invitrogen, cod. P36931) and examined by confocal fluorescence microscopy.

### SARS-CoV-2 propagation and infection

SARS-CoV-2 was isolated from a nasopharyngeal swab retrieved from a patient with COVID-19 (Caucasian man of Italian origin, genome sequences available at GenBank (SARS-CoV-2-UNIBS-AP66: ERR4145453)). This SARS-CoV-2 isolate clustered in the B1 clade, similar to most Italian sequences, together with sequences derived from other European countries and the United States. SARS-CoV-2 inoculum (a kind gift of Professor Arnaldo Caruso, University of Brescia) was obtained in Vero E6 cells. As previously described, viral titer was determined by plaque assay in Vero E6 cells^30^. SARS-CoV-2 manipulation was performed in the BSL-3 laboratory of the University of Ferrara, following the biosafety requirements. Both Calu-3 and CFBE cell susceptibility to SARS-CoV-2 infection was assayed by infecting single cells with an MOI of 0.1 for 2 h at 37°C, as previously reported (approx. 2×10^5^ infectious virus particles per well)^30^. Twenty-four, 48 and 72 hours after infection, the infected cells were collected.

### Viral RNA detection

RNA extraction was performed 24, 48 and 72 hours post infection (hpi) using a MagMAX Viral/Pathogen Nuclei Acid Isolation kit (Thermo Fisher, Italy) for recovery of RNA and DNA from the virus, as previously described^30^. SARS-CoV-2 titration was obtained using a TaqMan 2019nCoV assay kit v1 RealTime-PCR (Thermo Fisher, Italy).

### Cytokine assays

Well-differentiated primary HNECs were cultured in air-liquid interface (ALI) conditions and left unstimulated or primed with SARS-CoV-2 spike protein (1 μg/ml) on both the apical and basolateral sides to maximize the cellular response for 16 h at 37°C, and 5% CO_2_. IL-6 released into cell culture supernatants was measured by ELISA (ab46027, Abcam) following the manufacturer’s protocol. Samples were subsequently analyzed on a Sunrise microplate reader (Tecan Trading AG, Männedorf, Switzerland).

## Acknowledgements

We are grateful to the Cystic Fibrosis Foundation (Bethesda, MD) for kindly providing the CRISPR/Cas9 gene edited 16HBE14o-, W1282X-CFTR and G542X-CFTR cell lines, to Dr. J.R. Riordan (University of North Carolina, Chapel Hill) for anti-CFTR ab596 (Cystic Fibrosis Foundation Therapeutics), to Federica Quiri (Azienda Ospedaliera Universitaria Integrata, Verona, Italy) for the excellent technical support, and to Marc Chanson (University of Geneva) for providing Calu-3 SH3 and Alter cell lines. AR is supported by local founds from University of Ferrara, FIR-2021, the Italian Ministry of Health (GR-2016-02364602) and the Italian Ministry of Education, University and Research (PRIN Grant 2017XA5J5N). PP is supported by Italian Association for Cancer Research (AIRC, IG-23670), Telethon (GGP11139B), local funds from the University of Ferrara, and the Italian Ministry of Education, University and Research (PRIN Grant 2017E5L5P3). This work was partially funded by the following (alphabetically): Chiesi Farmaceutici (Parma, Italy), Mylan N.V. (Hatfield, UK), Pfizer Inc. (New York, NY). This work is part of the CF Italian Task Force Against COVID-19 Action (CF-ITACA) of the Italian Cystic Fibrosis Society (SIFC), which partially supported this work as well.

## Author contributions

VB conceived the idea, co-designed the study, analyzed the data, and wrote the manuscript; VG performed the in vitro viral infections and analyzed the data; MA, DO, MD, ET performed experiments concerning ACE2, CD13 and CFTR expression and ELISA; AF and CP performed the experiments with pre-miR-145; SL performed the IF experiments; AT, SV and GC analyzed the data and internally revised the manuscript; MB designed and performed the experiments on cytokine expression; PP, FB and GL critically reviewed the final manuscript and gave the final approval of the version to be published; RG designed the pre-miR-145 experiments and internally reviewed the manuscript; AR designed the IF experiments, analyzed the data, and internally reviewed the manuscript; RR designed and supervised the experiments concerning in vitro viral infections; MC conceived the idea, co-designed and supervised the study.

## References

1. Harrison, S. L., Fazio-Eynullayeva, E., Lane, D. A., Underhill, P. & Lip, G. Y. H. Comorbidities associated with mortality in 31,461 adults with COVID-19 in the United States: a federated electronic medical record analysis. PLoS Med. 17, e1003321 (2020).

2. Berardis, S. et al. SARS-CoV-2 seroprevalence in a Belgian cohort of patients with cystic fibrosis. J. Cyst. Fibros. 19, 872–874 (2020).

3. Bezzerri, V., Lucca, F., Volpi, S. & Cipolli, M. Does cystic fibrosis constitute an advantage in COVID-19 infection? Ital. J. Pediatr. 46, 143 (2020).

4. Mondejar-Lopez, P. et al. Impact of SARS-CoV-2 infection in patients with cystic fibrosis in Spain: incidence and results of the national CF-COVID19-Spain survey. Respir. Med. 170, 106062 (2020).

5. Corvol, H. et al. First wave of COVID-19 in French patients with cystic fibrosis. J. Clin. Med. 9, 3624 (2020).

6. Hoffmann, M. et al. SARS-CoV-2 cell entry depends on ACE2 and TMPRSS2 and Is blocked by a clinically proven protease inhibitor. Cell 181, 271–280.e8 (2020).

7. Walls, A. C. et al. Structure, function, and antigenicity of the SARS-CoV-2 spike glycoprotein. Cell 181, 281–292.e6 (2020).

8. Yan, R. et al. Structural basis for the recognition of SARS-CoV-2 by full-length human ACE2. Science 367, 1444–1448 (2020).

9. Luo, M. et al. IL-6 and CD8+ T cell counts combined are an early predictor of in-hospital mortality of patients with COVID-19. JCI Insight 5, e139024 (2020).

10. Cummings, M. J. et al. Epidemiology, clinical course, and outcomes of critically ill adults with COVID-19 in New York city: a prospective cohort study. Lancet 395, 1763–1770 (2020).

11. Qi, F., Qian, S., Zhang, S. & Zhang, Z. Single cell RNA sequencing of 13 human tissues identify cell types and receptors of human coronaviruses. Biochem. Biophys. Res. Commun. 526, 135–140 (2020).

12. Devarakonda, C. K. V., Meredith, E., Ghosh, M. & Shapiro, L. H. Coronavirus receptors as immune modulators. J. Immunol. 206, 923–929 (2021).

13. Chomarat, P., Rissoan, M. C., Pin, J. J., Banchereau, J. & Miossec, P. Contribution of IL-1, CD14, and CD13 in the increased IL-6 production induced by in vitro monocyte-synoviocyte interactions. J. Immunol. 155, 3645–3652 (1995).

14. Liang, W. et al. Possible contribution of aminopeptidase N (APN/CD13) to migration and invasion of human osteosarcoma cell lines. Int. J. Oncol. 45, 2475–2485 (2014).

15. Zotz, J. S. et al. CD13/aminopeptidase N is a negative regulator of mast cell activation. FASEB J. 30, 2225–2235 (2016).

16. Cohen-Cymberknoh, M., Kerem, E., Ferkol, T. & Elizur, A. Airway inflammation in cystic fibrosis: molecular mechanisms and clinical implications. Thorax 68, 1157–1162 (2013).

17. Viviani, L., Assael, B. M., Kerem, E. & ECFS (A) H1N1 Study Group. Impact of the a (H1N1) pandemic influenza (season 2009-2010) on patients with cystic fibrosis. J. Cyst. Fibros. 10, 370–376 (2011).

18. Somayaji, R. et al. Cystic fibrosis pulmonary exacerbations attributable to respiratory syncytial virus and influenza: a population-based study. Clin. Infect. Dis. 64, 1760–1767 (2017).

19. Colombo, C. et al. SARS-CoV-2 infection in cystic fibrosis: a multicentre prospective study with a control group, Italy, February-July 2020. PLoS One 16, e0251527 (2021).

20. Naehrlich, L. et al. Incidence of SARS-CoV-2 in people with cystic fibrosis in Europe between February and June 2020. J. Cyst. Fibros. 20, 566–577 (2021).

21. Braga, L. et al. Drugs that inhibit TMEM16 proteins block SARS-CoV-2 spike-induced syncytia. Nature 594, 88–93 (2021).

22. Chang, M. H. et al. Slc26a9 is inhibited by the R-region of the cystic fibrosis transmembrane conductance regulator via the STAS domain. J. Biol. Chem. 284, 28306–28318 (2009).

23. Kunzelmann, K., Kiser, G. L., Schreiber, R. & Riordan, J. R. Inhibition of epithelial Na+ currents by intracellular domains of the cystic fibrosis transmembrane conductance regulator. FEBS Lett. 400, 341–344 (1997).

24. Konstas, A. A., Koch, J. P., Tucker, S. J. & Korbmacher, C. Cystic fibrosis transmembrane conductance regulator-dependent up-regulation of Kir1.1 (ROMK) renal K+ channels by the epithelial sodium channel. J. Biol. Chem. 277, 25377–25384 (2002).

25. Riquelme, S. A. et al. Cystic fibrosis transmembrane conductance regulator attaches tumor suppressor PTEN to the membrane and promotes anti Pseudomonas aeruginosa immunity. Immunity 47, 1169–1181.e7 (2017).

26. Watson, M. J. et al. The cystic fibrosis transmembrane conductance regulator (CFTR) uses its C-terminus to regulate the A2B adenosine receptor. Sci. Rep. 6, 27390 (2016).

27. Scheckenbach, K. E. et al. Prostaglandin E²regulation of cystic fibrosis transmembrane conductance regulator activity and airway surface liquid volume requires gap junctional communication. Am. J. Respir. Cell Mol. Biol. 44, 74–82 (2011).

28. Ma, T. et al. Thiazolidinone CFTR inhibitor identified by high-throughput screening blocks cholera toxin-induced intestinal fluid secretion. J. Clin. Invest. 110, 1651–1658 (2002).

29. Short, D. B. et al. An apical PDZ protein anchors the cystic fibrosis transmembrane conductance regulator to the cytoskeleton. J. Biol. Chem. 273, 19797–19801 (1998).

30. Bortolotti, D. et al. TLR3 and TLR7 RNA sensor activation during SARS-CoV-2 infection. Microorganisms 9, 1820 (2021).

31. Fabbri, E. et al. A peptide nucleic acid against MicroRNA miR-145-5p enhances the expression of the cystic fibrosis transmembrane conductance regulator (CFTR) in calu-3 cells. Molecules 23, 71 (2017).

32. Sultan, S. et al. A peptide nucleic acid (PNA) masking the miR-145-5p binding site of the 3’UTR of the cystic fibrosis transmembrane conductance regulator (CFTR) mRNA enhances CFTR expression in calu-3 cells. Molecules 25, 1677 (2020).

33. Tamanini, A. et al. A peptide-nucleic acid targeting miR-335-5p enhances expression of cystic fibrosis transmembrane conductance regulator (CFTR) gene with the possible involvement of the CFTR scaffolding protein NHERF1. Biomedicines 9, 117 (2021).

34. De Santi, C. et al. Precise targeting of miRNA sites restores CFTR activity in CF bronchial epithelial cells. Mol. Ther. 28, 1190–1199 (2020).

35. Oglesby, I. K., Chotirmall, S. H., McElvaney, N. G. & Greene, C. M. Regulation of cystic fibrosis transmembrane conductance regulator by microRNA-145, -223, and -494 is altered in ΔF508 cystic fibrosis airway epithelium. J. Immunol. 190, 3354–3362 (2013).

36. Yang, X. et al. High CFTR expression in Philadelphia chromosome-positive acute leukemia protects and maintains continuous activation of BCR-ABL and related signaling pathways in combination with PP2A. Oncotarget 8, 24437–24448 (2017).

37. Roesch, E. A., Nichols, D. P. & Chmiel, J. F. Inflammation in cystic fibrosis: an update. Pediatr. Pulmonol. 53, S30–S50 (2018).

38. Majka, G. et al. Chronic bacterial pulmonary infections in advanced cystic fibrosis differently affect the level of sputum neutrophil elastase, IL-8 and IL-6. Clin. Exp. Immunol. 205, 391–405 (2021).

39. McGreal, E. P. et al. Inactivation of IL-6 and soluble IL-6 receptor by neutrophil derived serine proteases in cystic fibrosis. Biochim. Biophys. Acta 1802, 649–658 (2010).

40. Colombo, C. et al. Cytokine levels in sputum of cystic fibrosis patients before and after antibiotic therapy. Pediatr. Pulmonol. 40, 15–21 (2005).

41. Recchiuti, A. et al. Resolvin D1 and D2 reduce SARS-CoV-2-induced inflammatory responses in cystic fibrosis macrophages. FASEB J. 35, e21441 (2021).

42. Rimessi, A. et al. Mitochondrial Ca2+-dependent NLRP3 activation exacerbates the Pseudomonas aeruginosa-driven inflammatory response in cystic fibrosis. Nat. Commun. 6, 6201 (2015).

43. Bebok, Z. et al. Failure of cAMP agonists to activate rescued deltaF508 CFTR in CFBE41o-airway epithelial monolayers. J. Physiol. 569, 601–615 (2005).

