## Supplemental Table 1 for "ACE2 expression and localization are regulated by CFTR: implications beyond cystic fibrosis"

| Cell type | Sample ID | Gender | Age | Genetics (CFTR) |
|---|---|---|---|---|
| **CF** | | | | |
| HNEC | CF-MD0408 | N/A | 24 | Homozygote F508del |
| HNEC | CF-803 | N/A | 25 | Homozygote F508del |
| HBEC | CF-502 | M | 25 | Homozygote F508del |
| HBEC | CF-MD067301 | F | 23 | Homozygote F508del |
| HBEC | CF-MD056701 | F | 39 | Homozygote F508del |
| HBEC | CF- MD067901 | F | 30 | Homozygote F508del |
| **Control** | | | | |
| HNEC | MP0009 | N/A | pool* | normal |
| HBEC | MP076801 | F | 59 | normal |
| HNEC | MD0742 | M | 61 | normal |
| HBEC | MD064201 | F | 21 | normal |
| HBEC | MD067001 | M | 15 | normal |

**Supplementary Table 1. Genetics and clinical data of the human samples analyzed in this study.** N/A, not available; F, female; * pool of HNEC isolated from 14 heathy donors
